# G1 and G2 ApolipoproteinL1 modulate macrophage inflammation and lipid accumulation through the polyamine pathway

**DOI:** 10.1101/2025.06.06.658371

**Authors:** Esther Liu, Matthew Wright, Andrew O. Kearney, Tiffany Caza, Johnson Y. Yang, Valerie Garcia, Amal O. Dadi, Shuta Ishibe, Navdeep S. Chandel, Hanrui Zhang, Edward B. Thorp, Jennie Lin

## Abstract

The G1 and G2 variants of the gene encoding Apolipoprotein L1 (*APOL1*) increase risk for kidney disease and cardiometabolic traits. While previous studies have elucidated key mechanisms by which G1 and G2 *APOL1* cause cellular inflammation and cytotoxicity, it remains unclear whether these mechanisms drive inflammation in G1 and G2 macrophages. In this study, we used mouse bone-marrow-derived macrophages and human induced pluripotent stem cell-derived macrophages to identify altered immune signaling and inflammatory activation caused by G1 and G2 *APOL1*. We demonstrated that G1 and G2 APOL1 increased lipid accumulation, pro-inflammatory cytokine expression, and inflammasome signaling; this inflammatory response was sustained when treated with anti-inflammatory cytokines IL-4 and IL-10. Additionally, in G1 and G2 macrophages we observed increased mitochondrial size and elongation, oxidative phosphorylation, and glycolysis. Finally, we used unbiased metabolite analysis to identify an accumulation of polyamine spermidine and the enrichment of the spermidine synthesis pathway in G1 and G2 macrophages. When treated with polyamine inhibitor α-difluoromethylornithine (DFMO), lipid accumulation and inflammasome gene expression decreased in G1 and G2 macrophages. Together, these findings establish the pro-inflammatory effects of G1 and G2 *APOL1* in macrophages and identify a novel pathway which ameliorates G1 and G2 effects on cellular inflammation.

## Introduction

Chronic kidney disease (CKD) affects approximately 10% of the global population, and more than 14% of the United States population (1, 2). Within CKD, there is a disparity of greater disease prevalence among the US African American population (3). Beyond kidney disease, African American individuals are also more at risk of developing atherosclerosis, obesity, and hypertension (4–6). Part of this disparity is attributed to two risk alleles within the gene encoding Apolipoprotein L1 (*APOL1*) termed G1 and G2 (7, 8). The G1 variant contains the non-synonymous coding variants rs73885319 (S342G) and rs60910145 (I384M). The G2 contains the 6 base pair deletion (rs71785313) which deletes two amino acids N388 and Y389. Individuals with 2 copies of the *APOL1* G1 or APOL1 G2 risk variants (RV) face increased risk for kidney disease and possibly cardiometabolic traits compared to those with at least one copy of the non-risk allele (G0). A recent study also demonstrated that carrying one *APOL1* risk allele can increase the risk of CKD in West Africans (9). Numerous reports confirm that *APOL1* risk alleles increase the risk for focal segmental glomerulosclerosis, HIV-associated nephropathy, and COVID-19-associated nephropathy, all of which fall within the spectrum of APOL1-mediated kidney disease (AMKD) (7, 10–12). Some reports have also shown an association between *APOL1* risk alleles and atherosclerosis, obesity, and hypertension (13–15). The specific molecular mechanisms underlying this increased risk are still under investigation.

G1 and G2 APOL1 arose due to evolutionary pressure against trypanosomiasis and are only found in individuals with recent African ancestry. APOL1 evolved as a resistance factor to trypanosomes which cause African trypanosomiasis, *Trypanosoma brucei rhodesiense* and *Trypanosoma brucei gambiense* (16). APOL1 G1 and G2 variants are able to escape the trypanosome’s serum resistance-associated (SRA) proteins and create a pore in the trypanosome lysosomal compartments, leading to lysis (17). This pore-forming function of APOL1 has been shown to increase cation flux in epithelial cells leading to cytotoxicity (18). A new small molecule inhibitor of APOL1, VX-147, targets this pore-forming function in the context of podocytopathy and has shown some efficacy against AMKD in a phase 2a clinical trial (19, 20). However, it is not known whether this channel activity occurs in immune cells and if VX-147 would be efficacious in those cell types.

During infection, the trypanosome faces the challenge of evading the host’s immune system, which includes trypanolytic factors such as APOL1 but also innate immune responses from macrophages, which phagocytose the trypanosome (21). In particular, trypanosomes secrete metabolites including indole-3-pyruvate to modulate metabolic signaling and inhibit the macrophage’s ability to mount a pro-inflammatory response to kill the trypanosome (22, 23). During trypanosome infection, it is advantageous for the macrophage to have an increased pro-inflammatory response to counteract immunosuppressive signals from the trypanosome (22). However, outside of an infection context, sustained macrophage inflammation can lead to tissue injury.

As key regulators of tissue homeostasis, macrophages orchestrate responses to injury and inflammation through multiple functions including secreting cytokines, phagocytosing foreign bodies, and interacting with other cells in the environment (24, 25). When macrophage signaling is dysfunctional, this can lead to sustained inflammation and subsequent tissue fibrosis, thus promoting the development of CKD and other cardiometabolic traits characterized by low-grade chronic inflammation (26, 27). While these complex diseases are linked to *APOL1* risk alleles and also involve macrophage dysfunction in their pathophysiology, it is not well understood how macrophages contribute to AMKD or possibly APOL1-mediated cardiometabolic disease. G1 and G2 APOL1 have demonstrated cytotoxic effects in epithelial cell models by dysregulated ion flux, enhancing ER stress, mitochondrial dysfunction, and inflammasome activation (20, 28–31). However, the effects of APOL1 in immune cells, e.g., macrophages, are relatively unknown. APOL1 is expressed in macrophages, and cellular stress in macrophages is directly related to inflammatory function and increasing inflammatory injury.

In this study, we explore the effects of G1 and G2 APOL1 in macrophage inflammatory signaling. Current literature has shown that macrophages expressing *APOL1* risk alleles have decreased maturation (32) and cholesterol efflux (33). However, the underlying cellular pathways driving these phenotypes and any *APOL1* genotype-specific changes in macrophage inflammation and function have not been explored. We hypothesized that G1 and G2 APOL1 cause organelle stress and dysfunction, leading to a sustained inflammatory response in macrophages. To test this hypothesis, we used transgenic mice expressing APOL1 G0, G1, or G2 to generate macrophages from bone marrow. We discovered a sustained inflammatory phenotype in G1 and G2 BMDMs compared to G0 which was linked to elongated mitochondria, increased respiratory capacity, and accumulation of spermidine, a polyamine metabolite.

## Results

### G1 and G2 APOL1 potentiate lipid accumulation in macrophages

Relevant to macrophage function, previous studies have shown that risk-variant forms of APOL1 trigger intracellular stress and inflammation (34–36). Therefore, we hypothesized that macrophages expressing G1 and G2 APOL1 promote tissue injury by not entering a reparative state after pro-inflammatory activation. To evaluate the role of macrophages in AMKD, we examined kidney biopsies of individuals diagnosed with focal segmental glomerulosclerosis (FSGS) in the context of a high-risk *APOL1* genotype. Light microscopy revealed foamy macrophages in the glomeruli of a high-risk genotype allograft (Figure 1A). Foamy macrophages were also seen in the glomeruli of individuals with high-risk *APOL1* genotype and collapsing glomerulopathy (Figure 1B).

**Figure 1.**
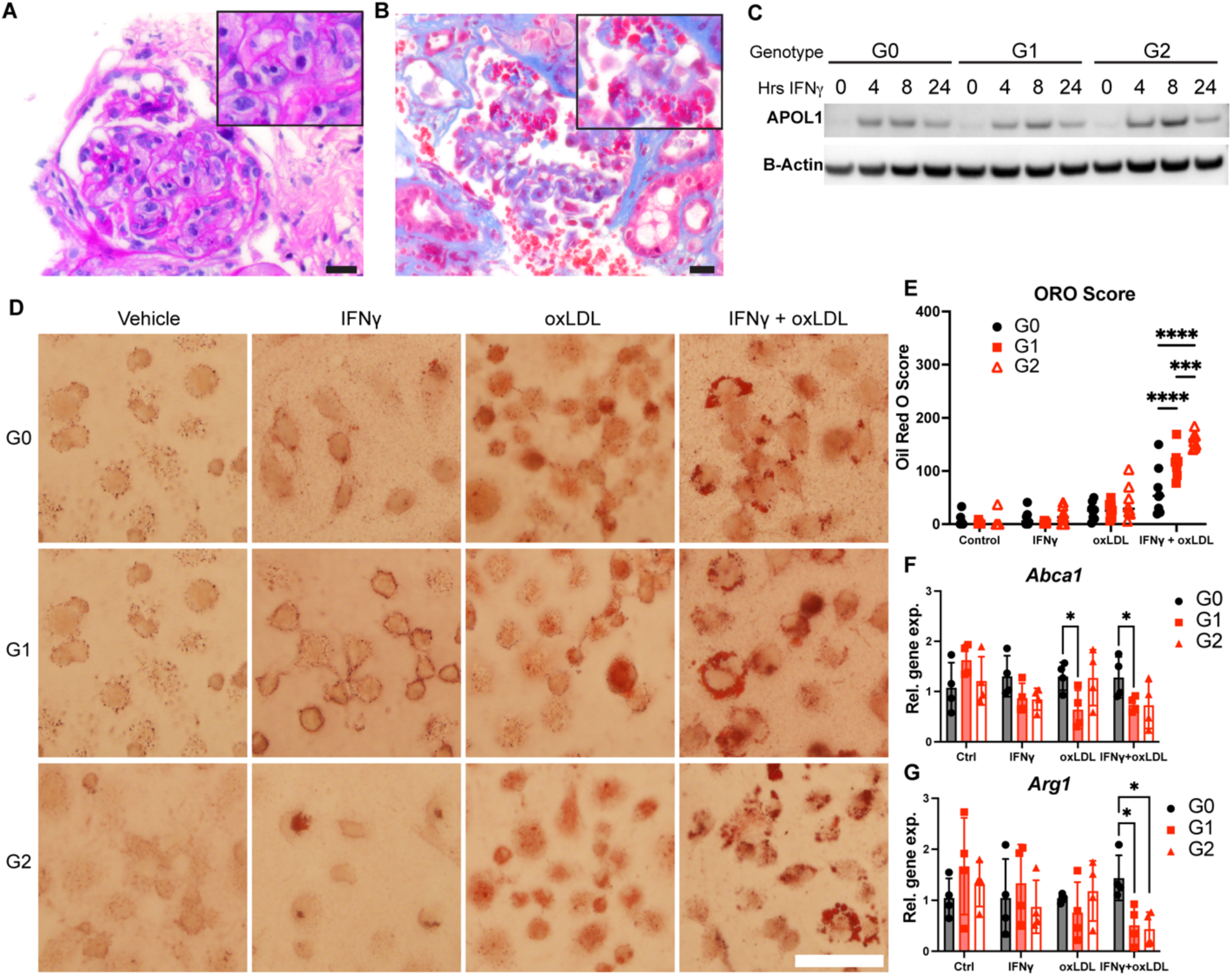
APOL1 genotype modulates lipid accumulation in macrophages. (**A**) Biopsy of kidney allograft from a donor with high-risk *APOL1* genotype reveals endocapillary foamy macrophages within the glomerulus (PAS, scale bar = 40 μm). (**B**) This glomerulus contains representative collapsing lesions seen in individuals with high-risk *APOL1* genotype. Endocapillary foamy macrophages are captured in this view (Masson’s trichrome, scale bar = 40 μm). (**C**) Western blot of APOL1 protein expression in BMDMs over a 24-hour time course of 5 ng/mL IFNγ treatment. (**D**) Oil Red O (ORO) stained images of BMDMs in the presence or absence of 5ng/mL IFNγ and 50 μg/mL oxidized LDL (oxLDL) for 72 hours. (**E)** Quantification of ORO staining. **(F**) Relative *Abca1* and (**G**) *Arg1* gene expression levels in BMDMs treated with IFNγ and oxLDL. Experiments were performed in BMDMs from 4-8 mice per genotype per group across 2 independent experiments, with both sexes represented. Data are expressed as mean ± SD. *p<0.05, **p<0.01 ***p<0.005 ****p>0.001. 2-way ANOVA with Tukey’s Multiple Comparison Test (**D**) and unpaired *t-test* (**G**).

The role of foamy macrophages in FSGS pathogenicity and progression has been unclear and is unknown in AMKD. To test the effects of *APOL1* genotype on foam cell formation, we analyzed lipid accumulation in bone marrow-derived macrophages (BMDMs) isolated from APOL1 G0, G1, and G2 transgenic mice. BMDMs from these mice expressed low levels of APOL1 at baseline and higher levels of APOL1 when treated with IFNγ, a known modifier of AMKD (Figure 1C; Supplemental Figure 1, A and B) (34, 37). We induced robust G0, G1, and G2 APOL1 expression in BMDMs with IFNγ and performed lipid loading with oxidized low-density lipoprotein (oxLDL) for 72 hours. Brightfield imaging of Oil Red O (ORO)-stained BMDMs revealed increased lipid accumulation in APOL1 G1 and G2 macrophages compared to G0 following incubation with IFNγ and oxLDL (Figure 1, D, and E). Similar results were seen in BMDMs harvested from a separate transgenic APOL1 mouse line and incubated with acetylated LDL (Supplemental Figure 2).

To investigate potential drivers of genotype-specific effects on macrophage lipid metabolism, we measured the expression of lipid transport genes. When treated with IFNγ and oxLDL, G1 and G2 BMDMs had significantly reduced expression of *Abca1*, a critical regulator of cholesterol efflux, potentially indicative of impaired lipid export (Figure 1F). Prior studies have shown that when macrophages become lipid-laden, they initially demonstrate anti-inflammatory patterns of gene expression but can develop phenotypic heterogeneity along the inflammatory spectrum (38–41). We hypothesized that lipid-loaded G1 and G2 macrophages would shift to a more pro-inflammatory state and found that *Arg1* expression, associated with a reparative macrophage phenotype, was significantly decreased in G1 and G2 foamy macrophages (Figure 1G). These findings suggest that G1 and G2 APOL1 contribute to dysregulated lipid metabolism in macrophages, potentially promoting foam cell formation and contributing to inflammatory disease processes.

### G1 and G2 APOL1 promote inflammatory signaling in macrophages

Because macrophage inflammation can promote foam cell formation (42, 43), we next investigated the effects of *APOL1* risk variants on macrophage activation. To directly test how *APOL1* variants alter inflammatory signaling, we treated murine and human macrophages carrying *APOL1* variants with either inflammatory or anti-inflammatory stimuli. First, we evaluated genotype-specific macrophage responses to induction of robust APOL1 expression with IFNγ. We observed relatively higher levels of pro-inflammatory markers in APOL1 G1 mouse BMDMs compared to G0, including *Tnf, Ccl2, Stat1, and Il18* on gene expression array and a trend towards higher CD86 expression on flow cytometry (Figure 2, A-D). APOL1 G2 BMDMs also expressed higher levels of *Cd86, Cd38,* and *Ccr2* compared to G0 but notably demonstrated a pro-inflammatory pattern that was more muted and distinct from G1 (Figure 2D). Upon further stimulation with LPS, G1 BMDMs demonstrated significantly higher CD86 expression (Figure 2, A-B). To validate the relevance of this result in a human model system, we tested the effects of IFNγ and LPS on genome-edited G0 and G1 human iPSC-derived macrophages (iPSDMs), which expressed relevant constitutive macrophage markers and APOL1 when treated with IFNγ (Supplemental Figure 1, C - E). G1 iPSDMs expressed higher levels of pro-inflammatory genes (Supplemental Figure 1F) while also secreting higher levels of pro-inflammatory cytokines, including IL-8, CXCL11, MIP-1α, and DPPIV (Figure 2E). Unexpectedly, despite mounting a more robust inflammatory response, macrophages expressing G1 APOL1 did not develop excess cytotoxicity previously reported in other cell types (20, 44, 45). Upon treatment with 24 hours of IFNγ, BMDMs demonstrated approximately 15% lower cell viability among all genotypes; G1 and G2 variants did not cause any additional cytotoxicity (Supplemental Figure 3).

**Figure 2.**
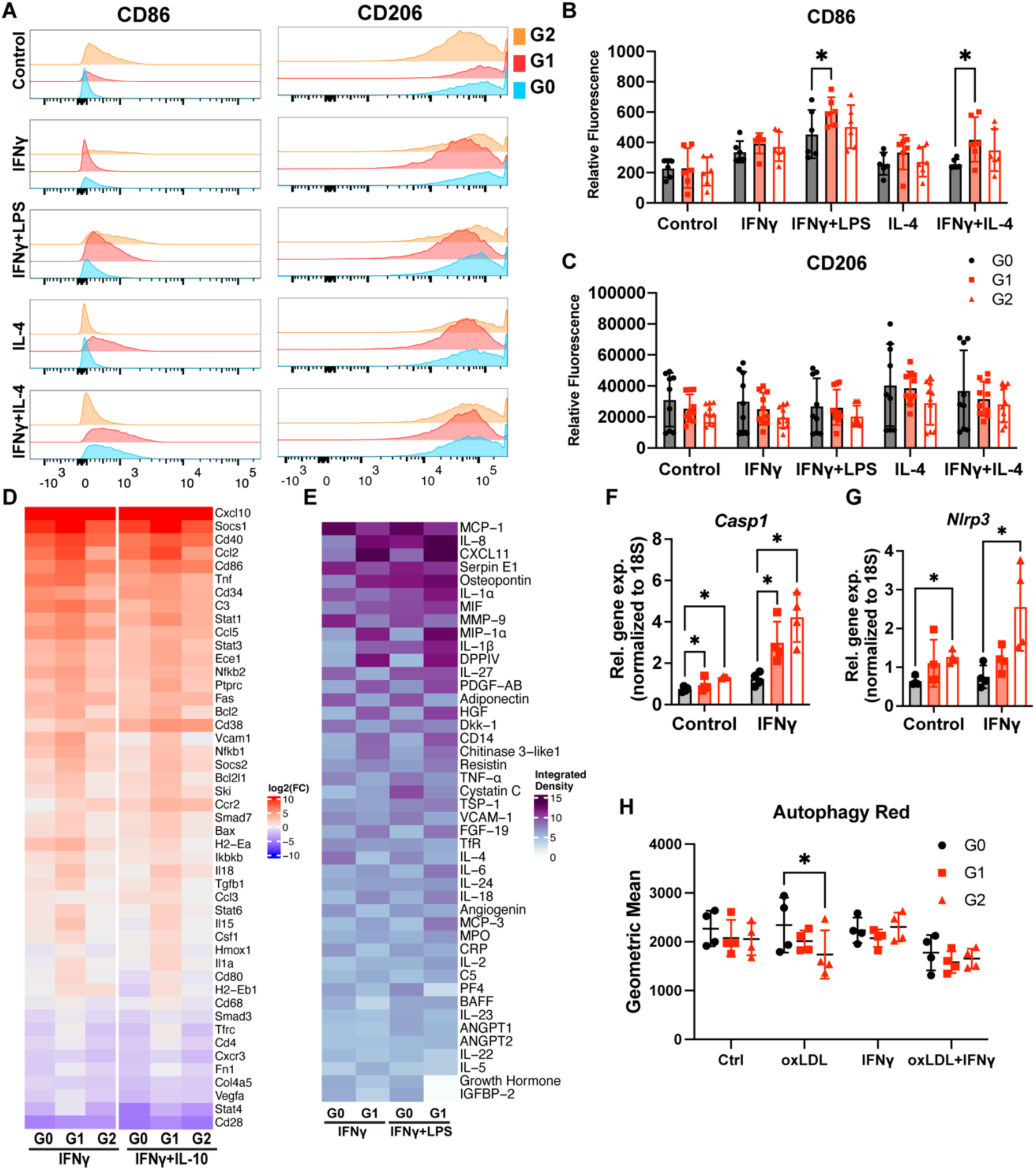
G1 APOL1 promotes higher levels of sustained inflammation. BMDMs treated with 5ng/mL IFNγ, 10 pg/mL LPS, and 10 ng/mL IL-4 for 8 hours were analyzed with flow cytometry to measure the surface expression of **(A)** CD86 and CD206. Histogram of **(B)** CD86 and **(C)** CD206 expression. **(D)** qPCR array of immune-related genes was completed with BMDMs treated with 5ng/mL IFNγ and 10 ng/mL IL-10. n=2 mice per genotype per treatment. **(E)** Dot blot measuring cytokines in iPSDM treated with 25ng/mL IFNγ and 10 pg/mL of LPS for 24 hours. Experiment was performed in n=2 iPSDM per genotype. **(F, G)** Gene expression of *Casp1, Nlrp3* in BMDMs treated with 5ng/mL IFNγ for 8 hours. **(H)** Autophagy measurement of BMDMs treated with IFNγ and oxLDL. Experiments were performed in BMDMs from 4-6 mice per genotype per group across 2 independent experiments, with both sexes represented **(A-C, F-G)**. Data are expressed as mean ± SD. 2-way ANOVA with Tukey’s Multiple Comparison Test (**B, H**) Unpaired t-test **(F-G)**, *p<0.05

In various chronic disease states, macrophage dysfunction involves failure to reprogram from a pro-inflammatory to a reparative state (46, 47). We hypothesized that G1 and G2 APOL1 promote sustained tissue inflammation by dysregulating macrophage response to anti-inflammatory signaling. Therefore, we tested whether the *APOL1* genotype modulates macrophage response to IL-4 and IL-10, which are known effectors in inflammation resolution (48, 49). After induction of APOL1 expression with IFNγ, IL-4 treatment resulted in higher expression of pro-inflammatory CD86 on flow cytometry in G1 BMDMs and higher transcript levels of *TNF* and *IL1B* in G1 iPSDMs (Figure 2, A and B; Supplemental Figure 1E), suggesting risk-variant APOL1 contributes to impaired reparative reprogramming. A similarly attenuated response to IL-10 was also seen in G1 BMDMs and for key genes such as *Cd86, Cd38, Fas,* and *Ccr2* in G2 BMDMs on the gene expression array (Figure 2D). These findings were not significantly different between male and female macrophages (Supplemental Figure 4).

Because inflammasome activation, a known inhibitor of reparative macrophage reprogramming (50), has been implicated in driving APOL1 podocytopathy (49), we next investigated whether inflammasome pathways are activated in G1 and G2 macrophages. We observed that in the presence and absence of IFNγ, *Casp1* expression was significantly upregulated in G1 and G2 BMDMs compared to G0 and that *Nrlp3* expression was significantly upregulated in G2 relative to G0 (Figure 2, F and G). Although G1 and G2 BMDMs had higher levels of NLRP3 protein than G0 as seen on Western blot, no genotype-specific differences in protein abundance were seen for AIM2, cleaved Caspase-1, and cleaved IL-1β (Supplemental Figure 5), suggesting that G1 and G2 macrophages are primed for increased inflammasome activity but that additional stressors are needed to activate the inflammasome.

In macrophages, activation of the inflammasome can be directly caused by defective autophagy (51), which has been observed in other APOL1 model systems (50). Additionally, defective autophagy can contribute to foam cell formation by impairing cholesterol efflux (52). Although we did not observe robust inflammasome activity in G1 and G2 macrophages stimulated with IFNγ, we hypothesized that G1 and G2 APOL1 would impair autophagic flux in macrophages, thereby promoting lipid accumulation seen in Figure 1D. To determine whether risk-variant forms of APOL1 disrupt macrophage autophagy, we performed a fluorescent autophagy assay in BMDMs. BMDMs incubated in the presence or absence of IFNγ and oxLDL for 72 hours did not demonstrate major genotype-specific differences in autophagic activity (Figure 2H), indicating that other mechanisms are driving G1 and G2 mediated dysregulation of macrophage inflammation and foam cell formation.

### G1 and G2 APOL1 localize to the mitochondria, altering mitochondrial shape and function in macrophages

To further investigate potential mechanisms by which G1 and G2 APOL1 contribute to dysregulated lipid metabolism and macrophage inflammation, we evaluated the localization of APOL1 within the macrophage. Depending on the model system and antibodies used, previous studies in epithelial cells have reported APOL1 localization at the plasma membrane (20, 53), mitochondria (54), and endoplasmic reticulum (ER) (55, 56). We observed, by immunostaining of BMDMs that after IFNγ induction, APOL1 protein colocalized with calnexin and TOM20, indicating that APOL1 is found at the ER and mitochondrial membranes, respectively (Figure 3A). This localization pattern was seen across the G0, G1, and G2 genotypes. Concordant with this finding, multiple groups previously reported involvement of ER stress in mechanisms of AMKD (29, 53, 57–59). However, markers of ER stress activation, including phospho-PERK, phospho-IRE1α, alternatively spliced *Xbp1*, and ATF4 were not significantly expressed in any APOL1 BMDMs regardless of genotype and IFNγ treatment (Supplemental Figure 6), establishing that ER stress is not a primary mechanism by which G1 and G2 APOL1 modulate macrophage function.

**Figure 3.**
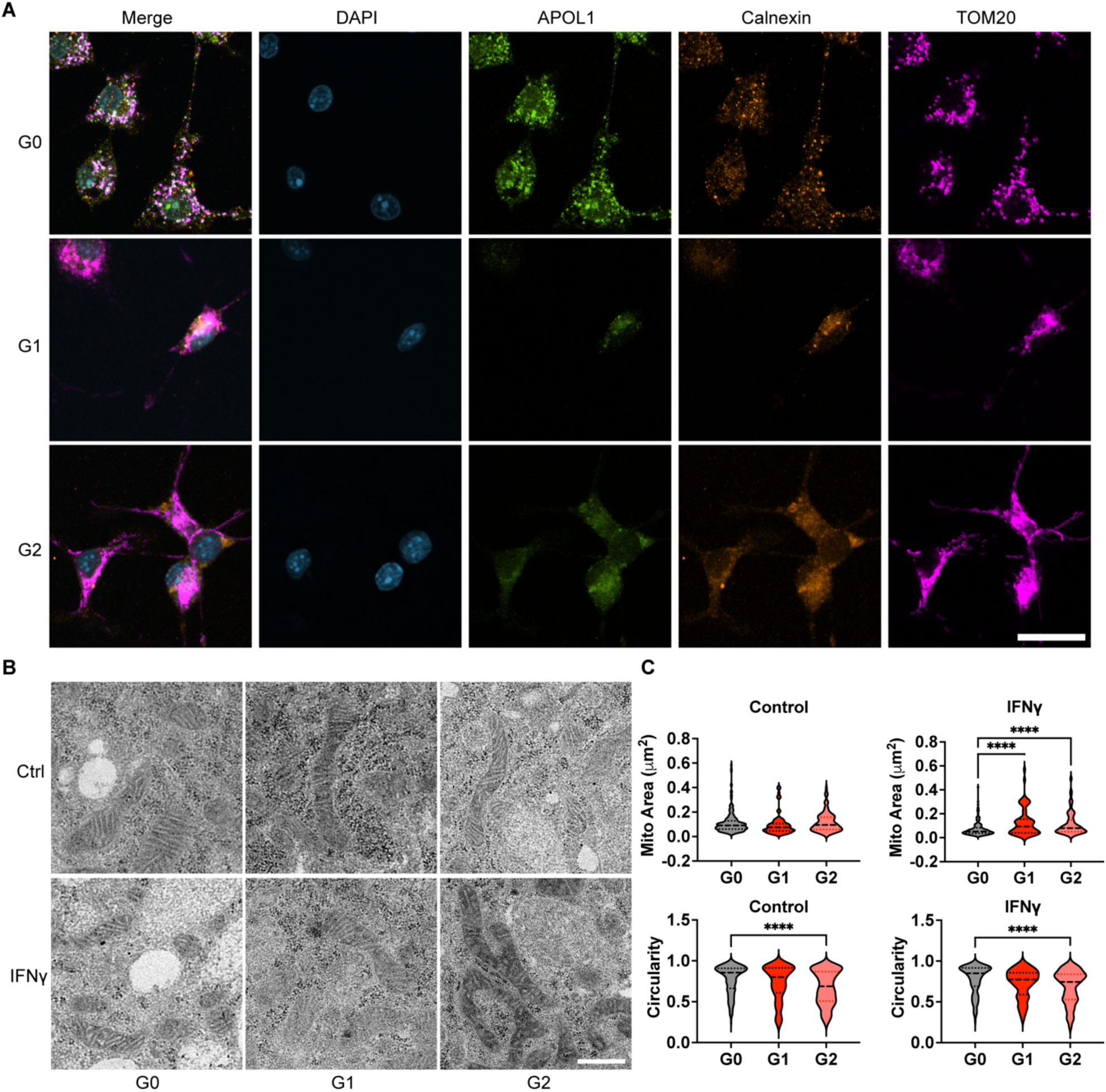
APOL1 localizes to the ER and mitochondria in macrophages and modulates mitochondrial morphology. **(A)** Representative images from confocal fluorescent microscopy of BMDMs stained with anti-APOL1, anti-Calnexin, and anti-Tom20 antibodies and DAPI. BMDMs were treated with 5ng/mL IFNγ for 8 hours. Experiment was performed in BMDMs from 4 mice per genotype per group. (Scale bar = 20 μm) **(B)** Representative transmission electron microscopy images of mitochondria in APOL1 BMDMs treated with 5 ng/mL IFNγ. (Scale bar = 500 nm) **(C)** Quantification of the area and circularity of BMDM mitochondria. Experiment was performed in BMDMs from 4 mice per genotype per group across 2 independent experiments, with both sexes represented. Measurements were performed in at least 100 mitochondria per sample. Data are expressed as median and interquartile ranges. Unpaired *t-test*, ****p>0.001

*APOL1* risk variants are linked to mitochondrial dysfunction, as shown in podocytes, but the effects in macrophages have not been determined. Because macrophage activation is linked to mitochondrial signaling, demonstrated by others to be dysregulated by risk-variant APOL1 (28, 54, 60–63), we also investigated G1 and G2 effects on macrophage mitochondria. Recently, mitochondrial dynamics between fission and fusion have been found to alter macrophage phenotype, with the initiation of reparative reprogramming requiring mitochondrial fission (64). To investigate the effects of G1 and G2 APOL1 on mitochondrial morphology in macrophages, we analyzed transmission electron microscopy (TEM) images of BMDMs expressing G0, G1, or G2 APOL1 (Figure 3B), quantifying mitochondrial area and circularity. Mitochondria in G1 and G2 macrophages exhibited increased area and reduced circularity compared to G0, indicating a shift toward elongated mitochondrial morphology consistent with more fusion seen in acute inflammation (Figure 3C). This finding was recapitulated in G1 iPSDMs (Supplemental Figure 7).

In addition to alterations in mitochondrial dynamics, increased glycolytic activity and oxidative phosphorylation (OXPHOS) are essential for pro-inflammatory macrophage activation. When we measured oxygen consumption rate (OCR) in our macrophage model system, G1 and G2 BMDMs demonstrated significantly elevated maximal and spare respiratory capacity when treated with IFNγ (Figure 4, A and B). Given this observed increase in OXPHOS, we next evaluated glycolytic capacity via proton efflux rate (PER) in IFNγ-treated G0, G1, and G2 BMDMs and found elevated basal glycolysis in IFNγ-treated G1 macrophages (Figure 4, C and D). Taken together, risk-variant forms of APOL1 in macrophages may promote sustained inflammation through local effects on mitochondrial morphology and downstream metabolism.

**Figure 4.**
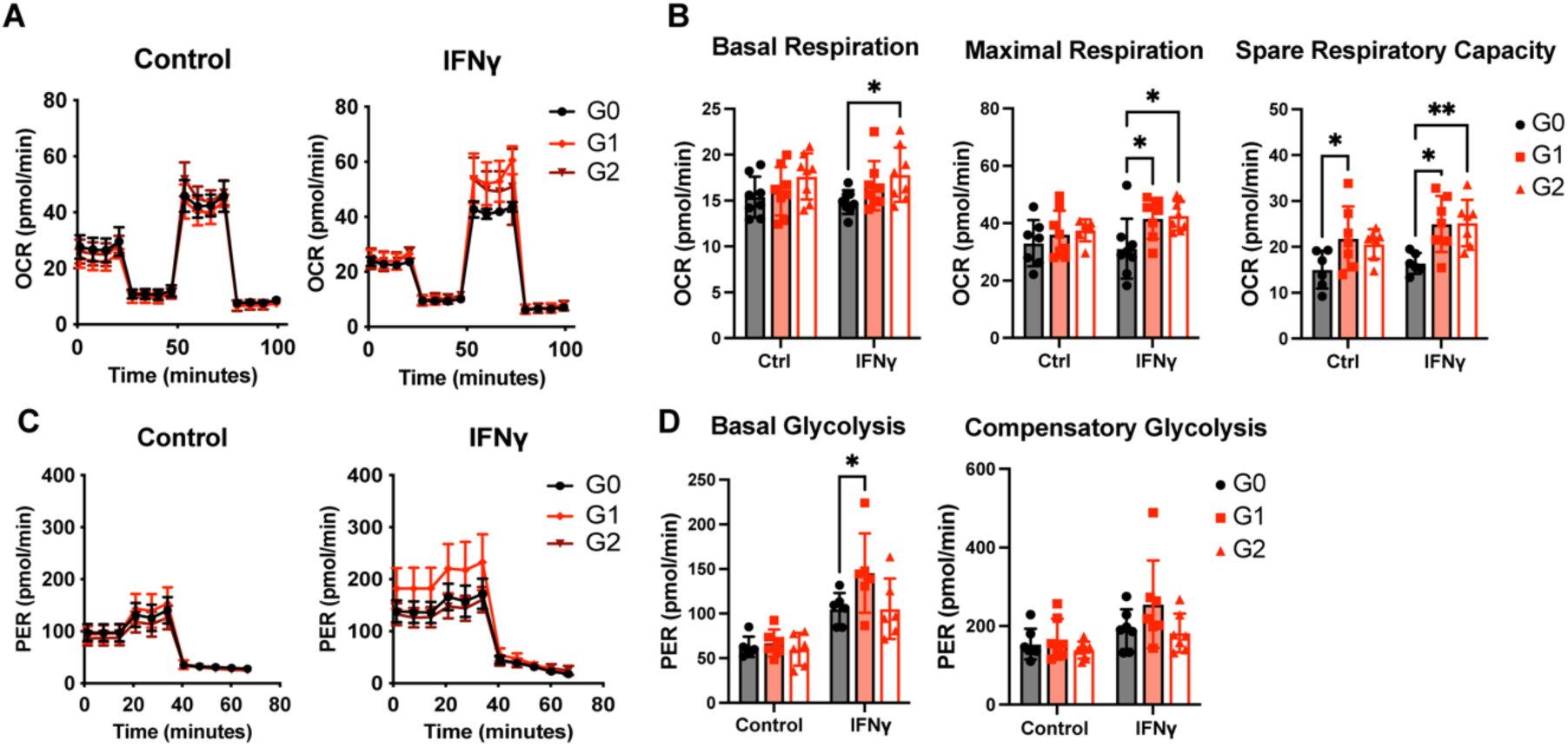
G1 and G2 APOL1 increase respiratory capacity. **(A)** Oxygen consumption rate of BMDMs treated with 5 ng/mL IFNγ or control measured by Seahorse Mito Stress Test. **(B)** Basal respiration, maximal respiration, and spare respiratory capacity of BMDMs from Mito Stress Test. **(C)** Proton efflux rate of BMDMs treated with 5 ng/mL IFNγ or control measured by Seahorse Glycolytic Rate Assay. **(D)** Basal and compensatory glycolysis measurement from Glycolytic Rate assay. Experiments were performed in BMDMs from 8 mice per genotype per group across 3 independent experiments, with both sexes represented. Data are expressed as mean ± SEM. 2-way ANOVA with Tukey’s Multiple Comparison Test, *p<0.05, **p<0.01

### G1 APOL1 mediates modest potassium efflux in macrophages

Notably, we did not observe APOL1 localization to the plasma membrane of BMDMs, despite the reported role of risk-variant APOL1 mediating cation transport that promotes osmotic swelling and cell death (45, 65, 66). Thus, APOL1-specific ion channel inhibitors such as VX-147 (19, 20, 67), currently in clinical trial, would potentially exert limited effects on macrophage phenotype and function. Because only a small portion of cell-intrinsic, risk-variant APOL1 localizes to the plasma membrane in epithelial cell models (20, 53), it is possible that the dynamic morphology could hinder effective visualization of plasma membrane colocalization by immunostaining in these macrophages. Therefore, to test the APOL1 ion channel hypothesis in macrophages at the functional level, we measured Thallium flux in G0, G1, and G2 BMDMs to evaluate genotype-specific effects on potassium transport. We found that when treated with IFNγ, G1 BMDMs demonstrated a modestly steeper Thallium flux slope than G0 BMDMs, a finding abrogated by the presence of VX-147 (Supplemental Figure 8), suggesting G1 APOL1 may potentially function as a cation channel on the plasma membrane. In contrast, no significant differences were found in ion flux for G2 BMDMs in the presence or absence of IFNγ and VX-147.

### Metabolomic analysis of APOL1 BMDMs reveals alternative polyamine regulation

To further investigate the effect of *APOL1* genotype on macrophage metabolic capacity, we performed steady-state metabolite analysis on APOL1 G0, G1, and G2 BMDMs. We identified the top differentially regulated metabolites in G1 and G2 BMDMs treated with IFNγ compared to G0 (Figure 5A, 5C). To our surprise, these metabolites were not involved in the TCA cycle. We identified pathways enriched in our data and quantified the “impact” of these pathways, calculated by giving more weight to central or rate-limiting metabolites. With an impact value of > 0.5, we identified 6 significant pathways, 5 of which are shared between G1 and G2 BMDMs (Figure 5B, 5D). The spermidine synthesis pathway involves the conversion of arginine into ornithine and the polyamine putrescine. Multiple metabolites relating to spermidine metabolism are significantly altered. Among the significantly altered metabolites, spermidine and N1/N8-acetylspermidine levels were significantly increased in G1 and G2 BMDMs relative to G0, while spermine and N1-acetylspermine were significantly decreased (Figure 5E). In G1 samples, N1-acetylspermine was not detected.

**Figure 5.**
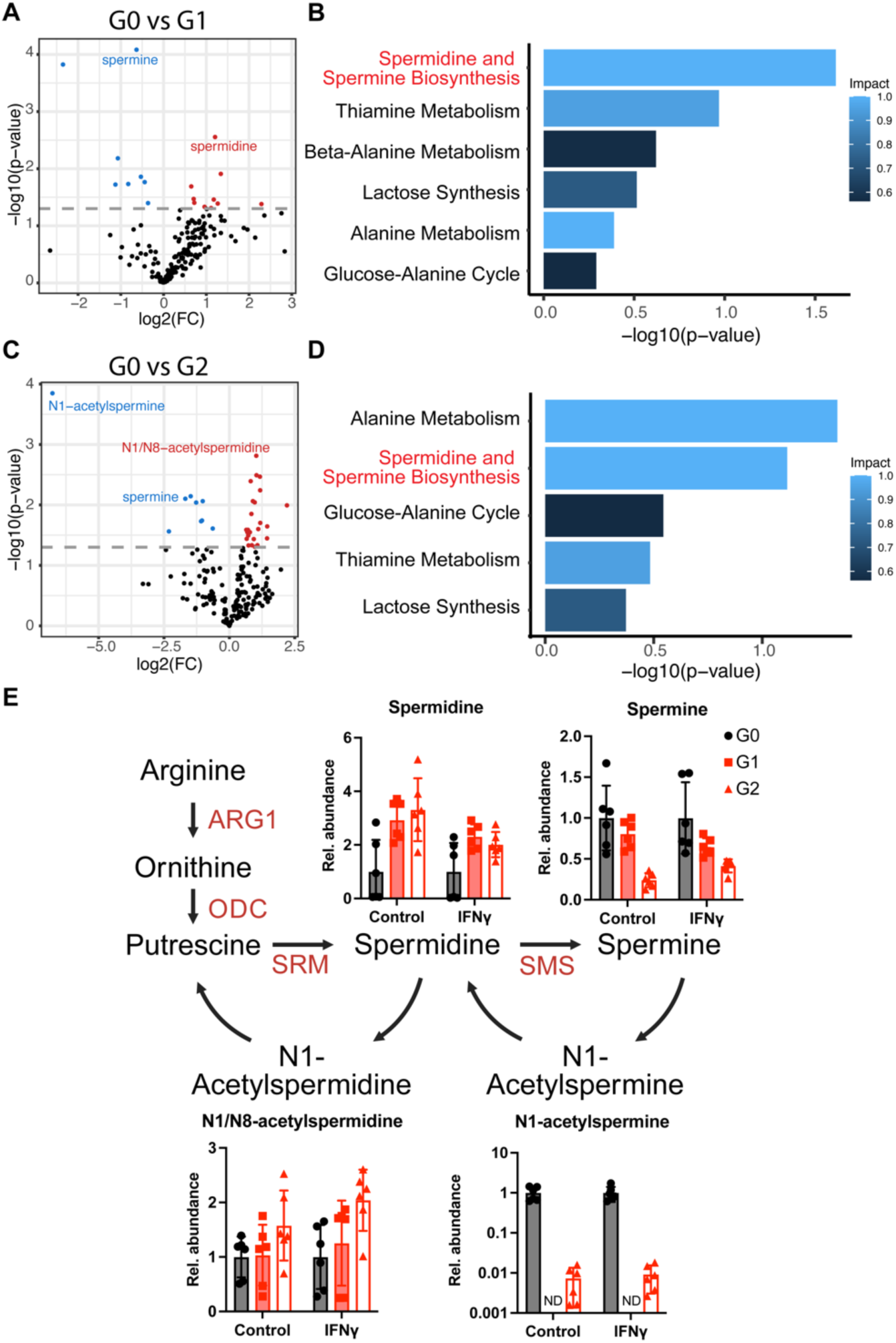
Metabolomics analysis of APOL1 BMDMS shows modified steady state of polyamines. **(A)** Volcano plot of significantly regulated metabolites in IFNγ treated G0 vs G1 BMDMs. **(B)** Most impactful pathways enriched in metabolomics analysis in G0 vs G1 BMDMs. **(C)** Volcano plot of significantly regulated metabolites in IFNγ treated G0 vs G2 BMDMs. **(D)** Most impactful pathways enriched in metabolomics analysis in G0 vs G2 BMDMs. **(E)** Fold change of metabolites in spermidine and spermine biosynthesis pathway. Experiments were performed in BMDMs from 3 mice per genotype per group across 2 independent experiments, with both sexes represented. ND = Not Detected

### Polyamine inhibitor α-difluoromethylornithine (DFMO) decreases lipid toxicity and inflammation in BMDMs

To investigate whether spermidine accumulation in G1 and G2 BMDMs contributes to their pro-inflammatory phenotype, we used polyamine inhibitor α-difluoromethylornithine (DFMO) to inhibit spermidine synthesis by ornithine decarboxylase (Figure 6A). BMDMs were treated with IFNγ, oxLDL, and DFMO for 72 hours and stained with ORO to visualize the accumulation of lipid droplets. The addition of DFMO alone did not alter ORO staining. However when treated with IFNγ, oxLDL, and DFMO, G1 and G2 BMDMs had significantly less ORO staining than G0 (Figure 6B, C). Additionally, expression of inflammasome genes (*Casp1*, *Nrlp3*) and pro-inflammatory cGAS-STING target genes (*Ccl2*, *Tnf*) was no longer significant between G0 and risk-variant APOL1 BMDMs except in G2 expression of *Tnf* (Figure 6D-G). These data suggest that reducing polyamine synthesis in macrophages can ameliorate the inflammatory phenotype caused by risk-variant APOL1.

**Figure 6.**
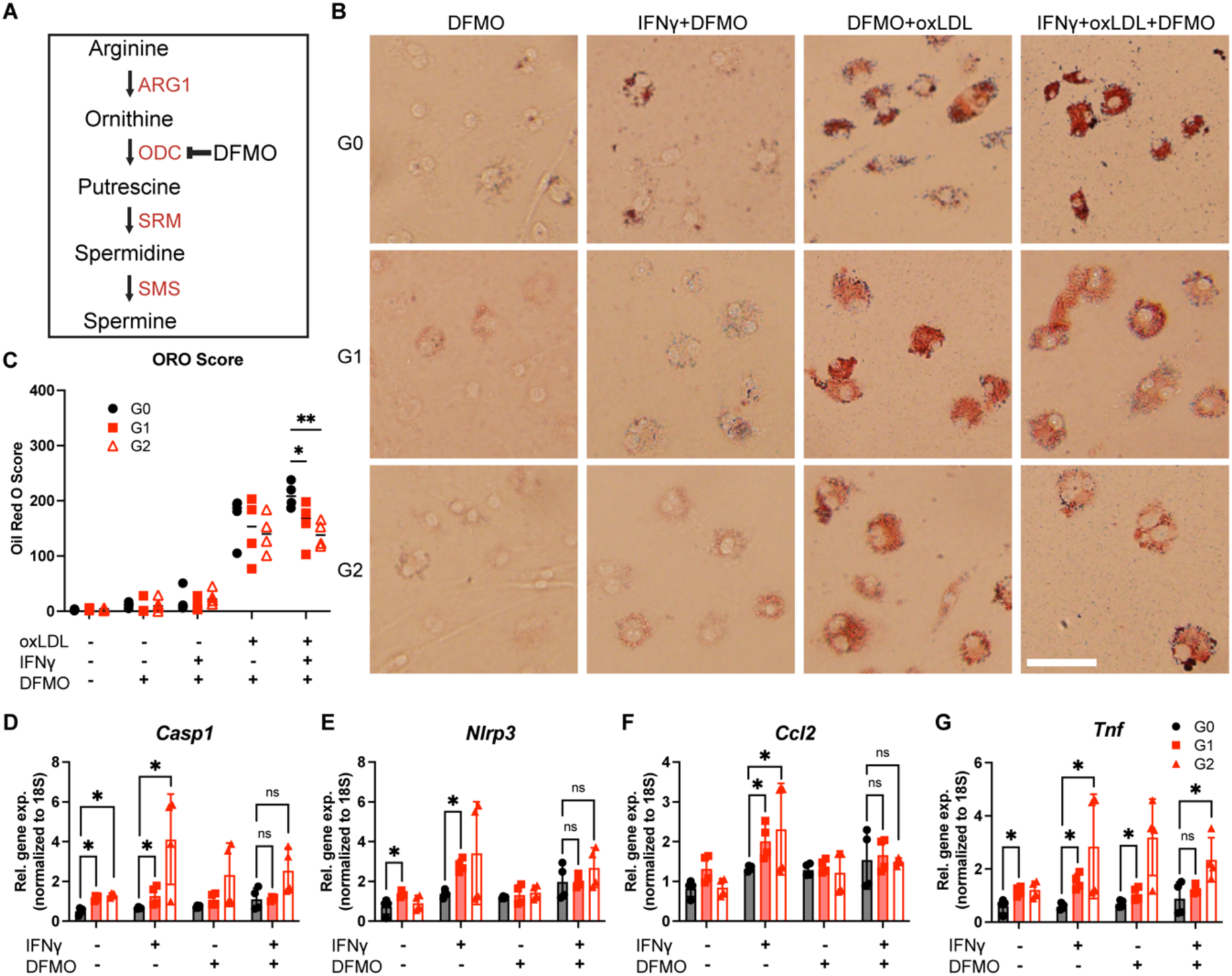
Polyamine inhibitor α-difluoromethylornithine (DFMO) decreases lipid toxicity and inflammation in BMDMs. **(A)** Mechanism of DFMO inhibition of polyamine synthesis pathway. **(B)** Representative images of BMDMs treated with IFNγ (5 ng/mL), oxLDL (50 μg/mL), and DFMO (200 μM) for 72 hours and stained with ORO. **(C)** Quantification of ORO stain. (Scale bar = 50 μm) **(D-G)** Gene expression of *Tnf, Ccl2, Casp1,* and *Nlrp3* in APOL1 BMDMs treated with IFNγ (5 ng/mL) and DFMO (200 μM) for 24 hours. Experiments were completed in BMDMS from 4 mice per genotype per group, with both sexes represented. Data are expressed as mean ± SD. Unpaired *t-test*, *p<0.05

## Discussion

APOL1’s role as an innate immunity factor has informed studies investigating mechanisms of podocyte injury (69, 70), but the role of its intrinsic expression in immune cells has not been as extensively explored. The APOL1 G1 and G2 variants are causal in multiple forms of kidney disease and are incompletely penetrant, leading to the “second-hit hypothesis” of immune system activation as a trigger for AMKD development (37, 71). The role of immune activation in AMKD has been seen in HIV-associated nephropathy, lupus nephritis, and COVID-19-associated nephropathy (14, 72–75), highlighting the potential impact of immune cell function dysregulated by risk-variant forms of APOL1. Additionally, G1 and G2 risk alleles have been associated with differences in cardiovascular traits, including obesity, stroke, and plaque rupture in coronary artery disease (76), all of which involve immune cells, and in particular the macrophage, in their development.

In the kidney and other tissues, macrophages maintain tissue homeostasis but can contribute to disease progression if they remain chronically inflamed (74). This study reveals novel findings of APOL1-mediated effects on macrophage function ultimately relevant to tissue injury and dysregulated repair. We observed higher levels of inflammation in G1 and G2 macrophages when robust APOL1 expression was induced by IFNγ, as well as more resistance to IL-4 and IL-10 signaling. Specifically, we observed increased expression of *Ccr2,* the gene for the receptor of monocyte chemoattractant protein-1 and increases monocyte-macrophage recruitment(77). This pro-inflammatory priming may also contribute to the formation of G1 and G2 foam cells, a nidus of chronic inflammation. Previous studies have shown that G1 and G2 macrophages have metabolic defects including decreased cholesterol efflux, maturation, and increased eicosanoid production (32, 33). Our oxLDL results confirmed Ryu et al’s findings of enhanced G1 and G2 lipid accumulation after aggregated LDL loading and ultimately connected these findings to metabolic dysfunction linked to the polyamine pathway (33).

Because prior studies revealed that risk-variant APOL1 can behave as a cytotoxic, druggable ion channel (61), we initially tested whether APOL1 genotype affected ion flux in macrophages. We observed a modest effect for G1 BMDMs that responded to VX-147. Because G1 and G2 BMDMs differ in their pro-inflammatory profiles, differential ion flux may play a role in G1-mediated macrophage dysfunction but would not explain G0 vs. G2 differences. Instead, we found that APOL1, regardless of genotype, colocalizes with the mitochondria and ER, so we hypothesized that G1 and G2 APOL1 amplify macrophage inflammation through organelle stress. The APOL1 protein structure is characterized to contain several functional domains including a membrane address domain which is necessary for its cytotoxic effects in 293T cells(79, 80). However, it is not clear whether these domains have any organelle or plasma membrane specificity. We had previously shown that G1 APOL1 promoted ER stress in kidney organoids (77) but, to our surprise, found no genotype-specific effects on ER stress in the macrophage.

Because macrophage activation is tightly regulated by metabolism and mitochondria (80), we evaluated mitochondrial dynamics and function, observing more mitochondrial elongation, higher oxygen consumption rate, and glycolytic activity in G1 and G2 macrophages compared to G0. The mitochondrial elongation in G1 and G2 BMDMs is not consistent with the increase in mitochondrial fission seen in primary G1/G1, G2/G2, and G1/G2 human proximal tubule cells treated with polyinosinic-polycytidylic acid in a prior study (81), a difference possibly attributable to cell type. Consistent with our data, mitochondria of macrophages treated with LPS have been noted to appear more elongated during the initial acute phase of inflammation and then undergo more fragmentation over time as the macrophages adopt a pro-resolving phenotype (81), highlighting cell-type specific mitochondrial dynamics. In addition, our data were consistent with more spare respiratory capacity in G2 and G2 macrophages compared to G0, which differs from prior findings in epithelial model systems (82). The relationship between OXPHOS and macrophage inflammation is complex and context-dependent (83–85), and it was unclear whether the genotype-specific differences in respiratory capacity were causal in differences in inflammatory response. To begin to investigate this, we performed metabolomics to identify perturbations of metabolites upstream and downstream of the TCA cycle. Although we had expected to find differential accumulation of TCA cycle metabolites relevant to APOL1’s localization, the significant finding from our metabolomics assay was enrichment in the polyamine pathway, which can modulate mitochondrial metabolism (86, 87).

While the effects of polyamines in macrophage activation are associated with increased autophagy and resolving inflammation, the overaccumulation of spermidine can result in cytotoxicity (88). Spermidine is also known to mediate the hypusination of translation factor EIF5A, which increases mitochondrial activity and fatty acid oxidation(89). Within the macrophage, the main source of polyamines comes from the conversion of L-arginine to L-ornithine which is subsequently decarboxylated into putrescine (90–92). Arginine itself is a key metabolite in the activation of macrophages, as it is used to generate nitric oxide during pro-inflammatory responses (93, 94). While arginine metabolism was not a significantly enriched pathway in our metabolic analyses, there remains more to understand about the connection between arginine metabolism, spermidine metabolism, and the pro-inflammatory effects of risk-variant APOL1 in macrophages.

Beyond its role in macrophage autophagy, polyamines have long been known to be a key metabolic target for trypanosomes, the same parasite against which APOL1 has evolved to provide protective immunity. Indeed, DFMO, also known as eflornithine, has been approved as a therapeutic against late-stage trypanosomiasis since 1990 and since then has acquired approvals for other indications including hirsutism and neuroblastoma (95). In trypanosomes, DFMO prevents the production of thiols used for the reduction of oxidative species and can prevent the production of variable surface glycoprotein, a necessary component of the cell surface (96). These connections between the macrophage, mitochondrial metabolism, and trypanosome metabolism necessitate further investigation.

Our study has several strengths, including (1) highlighting functional effects of APOL1 risk variants on macrophage phenotype and function relevant to cardiometabolic and kidney disease, (2) studies in both human and mouse model systems, and (3) identifying a novel pathway that attenuates G1 and G2 effects on inflammation and lipid metabolism in macrophages. Our work also has several limitations to consider. First, significant differences have been reported between mouse and human macrophages (97), so we leveraged a human-based iPSDM system to validate findings. Our results show that the inflammatory profile and mitochondrial morphology of G1 macrophages are consistent between human and mouse. Validation of results would be further bolstered if replicated in primary monocyte-derived macrophages from G0/G0 and AMKD patients. Second, macrophage function varies according to tissue microenvironment, and tissue-specific molecular data from *in vivo* experiments or single-cell molecular data from tissue biopsies would be informative in establishing the macrophage’s role in APOL1-mediated disease. These data would validate G1 and G2 macrophage effects in relevant disease models including kidney and cardiometabolic disease.

In conclusion, the risk-variant forms of APOL1 exert effects on macrophage function, promoting sustained inflammation and altering mitochondrial and lipid metabolism. These effects, which ultimately impact tissue homeostasis and repair, develop independently of APOL1’s role as an ion channel, indicating that mechanisms by which APOL1 alters cellular function are specific to cell type. Therapeutic targeting of the macrophage for enhanced treatment of AMKD or cardiometabolic traits would require novel strategies, such as addressing the polyamine pathway.

## Materials and Methods

### Induced pluripotent stem cell (iPSC)

iPSC lines previously derived from fibroblasts from a non-African ancestry donor (1016SevA; Harvard Stem Cell Institute) and from a healthy, Black, female donor (ATCC, BXS0114) were used in this study. Both cells lines are originally G0 *APOL1* genotype. CRISPR-Cas9 genome editing was performed to create G1 lines as previously described (29).

### APOL1 transgenic mice

APOL1 transgenic mice were obtained from the Mutant Mouse Resource & Research Centers at the University of California, Davis, and were developed as previously described (96). Briefly, bacterial artificial chromosome (BAC) containing the APOL1 gene as well as upstream (∼26kb) and downstream (∼65kb) regions were introduced into mice of FVB/NJ background. BACs containing the G0, G1, and G2 APOL1 variants were introduced, and mice for each genotype were generated. For additional validation studies, independently generated G0, G1, and G2 APOL1 BAC transgenic mice on a C57BL/6 background were a gift from Dr. Shuta Ishibe.

### Mouse husbandry and breeding

Transgenic APOL1 mice were housed in cages in a conventional animal facility at Northwestern University. Mice were housed at a maximum of five littermates per cage, with ad libitum food and H_2_O. All strains were maintained on a standard 12 h light cycle, at room temperature and humidity (20-22°C and 30-70%, respectively) as per the Guide for Care and Use of Laboratory Animals.

### Mouse bone marrow-derived macrophages

Bone marrow-derived macrophages (BMDMs) were generated as described in (97). In brief, bone marrow was harvested from femurs and tibias of mice 8-12 weeks of age and cultured in RPMI media supplemented with 10% heat-inactivated fetal bovine serum (Gibco, 10082147), 20% L929 cultured media, 1x GlutaMAX (Gibco 35050061), and 1% penicillin/streptomycin (10,000 U/mL, Gibco, 15140122). Media was replaced every 3 days and macrophages were used on days 7-10 of culture. Cells were cultured in a 37°C incubator with 5% CO_2_.

### iPSC culturing

iPSCs were cultured in a feeder-free system on 10 cm tissue culture dishes coated with Geltrex (Gibco, A1413201) substrate in mTeSR Plus (STEMCELL Technologies, 100-0276) supplemented with 0.02% Plasmocin (Invivogen, ant-mpp). iPSCs were cultured at 37° Celsius supplemented with 5% CO2. iPSCs were subcultured with 1/3 Accutase (STEMCELL Technologies, 07922) diluted in sterile 1X PBS (Gibco, 10010049). Cell lines were confirmed to be mycoplasma-free and below passage 55. Karyotyping of the iPSCs was completed and there were no chromosomal abnormalities.

### Generation of iPSC-derived macrophages (iPSDM)

iPSDM were generated by first using the STEMdiff Hematopoietic Kit (STEMCELL Technologies) to generate hematopoietic stem cells (HSC). HSCs were differentiated into macrophages following the protocol demonstrated by Yanagimachi et al.(98). Briefly, HSCs were cultured in StemPro-34 media supplemented with the following cytokines: Flt-3 ligand (50 ng/mL), GM-CSF (25 ng/mL) and M-CSF (50 ng/mL). The media was replaced every 3-4 days.

### Histological analysis of human kidney biopsies

Biopsies of kidneys with the high-risk *APOL1* genotype were examined at Arkana Laboratories under a waiver of consent. Samples shown were stained with periodic acid-schiff and Masson’s Trichrome stain. Images were taken with 40X objective (400x total magnification).

### Western blot

For whole cell lysate, iPSDMs and BMDMs were lysed with RIPA buffer (Sigma Aldrich, R0278) with protease and phosphatase inhibitor cocktail (Thermo Fisher Scientific, 78440) on ice for 10 minutes. Protein concentration was determined by Pierce’s BCA protein assay (Thermo Fisher Scientific, 23227). Samples were prepared using NuPage LDS sample buffer (Invitrogen, NP0007) with NuPAGE sample reducing agent (Invitrogen, NP0009) and heated for 10 minutes at 70° C. 15-20 µg protein samples were separated on a 4%-12% polyacrylamide NuPAGE Bis-Tris gel (Thermo Fisher Scientific, NP0321BOX). Gels were transferred to a PVDF membrane using the iBlot 2 transfer system (Invitrogen). Membranes were blocked with Superblock (TBS) blocking buffer (Thermo Fisher Scientific, 37535). Proteins were detected using anti-APOL1 (3.7D6, 1:1000, Genentech), anti-B-Actin (4970, 1:5000, Cell Signaling Technologies), anti-p-PERK (3197, 1:1000, Cell Signaling Technologies) anti-PERK (3192, 1:1000, Cell Signaling Technologies), anti-IRE1a (3294, 1:1000, Cell Signaling Technologies), anti-p-IRE1a (ab124945, 1:1000, Abcam), anti-ATF6 (65880, 1:1000, Cell Signaling Technologies), anti-NLRP3 (15101, 1:1000, Cell Signaling Technologies), anti-AIM2 (63660, 1:1000, Cell Signaling Technologies), anti-cleaved IL-1β (63124, 1:1000, Cell Signaling Technologies), anti-cleaved Caspase-1 (89332, 1:1000, Cell Signaling Technologies), and anti-ACS (67824, 1:1000, Cell Signaling Technologies) antibodies incubated overnight at 4°C with shaking. Secondary antibodies (Goat anti-rabbit IgG HRP-linked antibody, Ab6721, 1:2000, Abcam) conjugated with horseradish peroxidase were used and protein was detected using SuperSignal West Pico PLUS ECL substrate (Thermo Scientific, 34579) and imaged with an iBright CL1500 (Invitrogen). Densitometry was performed using the iBright Analysis Software (Invitrogen).

### Loading of oxidized low-density lipoprotein

BMDMs were seeded into gelatin-coated glass coverslips and pretreated with 5 ng/mL or 25 ng/mL of IFNγ or control for 8 hours. Media was then removed and replaced with macrophage serum-free media (Gibco, 12065074) containing 50ug/mL of oxidized low-density lipoprotein (oxLDL, Invitrogen, L34357) and incubated for 72 hours. Cells were then washed 3 times with PBS and fixed with 4% paraformaldehyde for 30 minutes at room temperature.

### Oil Red O staining

Fixed BMDMs were washed with 1x PBS for 5 minutes and 60% isopropanol for 15 seconds. A solution of 0.5% (%w/v) Oil Red O (Sigma Aldrich, O0625) in 60% isopropanol was prepared fresh before staining. Cells were stained with the Oil Red O solution for 1 minute. Cells were washed with water 3 times to remove excess stain. Cells were imaged with light microscopy. Images were scored for Oil Red O staining as previously described in Torous et al.(99) Briefly, 100 cells per sample were assigned a score of 0-4 based on the opacity of the cell with a score of “0” assigned to cells with no staining and “4” assigned to cells with a stain greater than ¾ opacity. The sum of the scores creates a scale of 0-400 to quantify the overall staining opacity of the sample. Samples were blinded during imaging and scoring to reduce bias during analysis.

### RNA isolation

RNA was isolated through TRIzol (Invitrogen, 15596026) and chloroform extraction and then purified with PureLink RNA Mini Kit (Invitrogen, 12183020) according to the manufacturer’s instructions. RNA quantity and quality were analyzed using a spectrophotometer (NanoDrop 2000, Thermo Fisher Scientific)

### Semi-quantitative RT-PCR

cDNA was generated using the High-Capacity cDNA Reverse Transcription Kit (Applied Biosystems, 4368814). *Xbp1* cDNA was amplified using BioRad Thermocycler and visualized using agarose gel electrophoresis. To amplify the spliced and unspliced *Xbp1* mRNA, *Xbp1* primers were used as described previously(100). PCR products were electrophoresed on 2.5% agarose gel. ACTB was used as a loading control. The size difference between the spliced and the unspliced *Xbp1* is 26 nucleotides.

### qRT-PCR

cDNA was generated using the High-Capacity cDNA Reverse Transcription Kit (Applied Biosystems, 4368814). Relative gene expression in the cDNA was analyzed with QuantStudio3 and QuantStudio5 (Thermo Fisher Scientific) using SYBR Green (Applied Biosystems, 4309155) or Taqman Universal PCR Master Mix (Applied Biosystems, 4444964). Each gene expression was normalized to 18S or *ACTB*. Primers for qRT-PCR are provided in the supplemental data.

### Macrophage polarization assay

Macrophages were treated with 5 ng/mL of recombinant mouse IFNγ (Fisher, 315-05), 10 pg/mL LPS (Sigma, L2630), or 10 ng/mL of IL-4 (Fisher, 200-04) for 8 hours to induce an M1 (IFNγ and LPS) or M2 (IL-4) phenotype. Macrophages were collected for downstream flow cytometric analyses.

### Flow cytometric analysis

Cells were dissociated from culture, washed with 1X PBS, and resuspended in flow buffer (1X PBS, 0.5% BSA, 0.05% Sodium Azide). The cells were resuspended in antibody solution at 1 × 10^!^cells/mL and incubated on ice in the dark for 30 minutes. The cells were washed with flow buffer twice and resuspended in PBS with Zombie Aqua viability stain (BioLegend, 423101) for 15 minutes at room temperature in the dark. Cells were washed with flow buffer three times and then incubated in an antibody cocktail containing fluorescently conjugated antibodies. Antibodies used include anti-CD14 (BD Biosciences, 561116), anti-CD16 (BD Biosciences, 556619), anti-CD64 (BD Biosciences, 560970), anti-CD11b (BD Biosciences, 561685), anti-Cd206 (Biolegend, 141720) and anti-Cd86 (Biolegend, 150613) antibodies. Cells were washed with flow buffer two times and resuspended in flow buffer for analysis on a Fortessa 1 analyzer (BD Biosciences).

### Immune response qPCR array

BMDMs were treated with 5 ng/mL of IFNγ and 10 ng/mL IL-10 or PBS control for 8 hours. RNA was isolated and cDNA was generated from the BMDMs. The TaqMan Mouse Immune Array v2.1 (no. 4365297, Thermo Fisher Scientific) was used to quantify 96 genes (including housekeeping controls) relating to immune response. The array was completed and analyzed with the QuantStudio™ 7 Flex Real-Time PCR System (Thermo Fisher Scientific).

### Cytokine dot blot

Supernatant from cultured iPSDMs treated 25 ng/mL IFNγ and 10 pg/mL LPS was collected and centrifuged at 300 x g for 10 minutes to remove any cellular debris. The cleared supernatant was analyzed using the Proteome Profiler Human XL Cytokine Array Kit (R&D systems) according to the manufacturer’s protocol. Membranes were imaged with iBright 1500 (Thermo Fisher Scientific).

### Immunofluorescence

Cells seeded on glass coverslips were fixed with 4% paraformaldehyde for 30 minutes at room temperature. After fixation, cells were blocked with 5% donkey serum (Abcam, ab7475) in a 1X PBS solution containing 0.3% Triton X-100 for 1 hour at room temperature. Cells were incubated with primary antibodies in a 1X PBS solution with 3% BSA and 0.3% Triton X-100 overnight at 4°C. Cells were washed three times with 1x PBS and stained with secondary antibodies for 1 hour at room temperature. Cells were washed and mounted onto slides for fluorescent microscopy with ProLong Gold antifade mountant (Invitrogen, P36930). Images were captured using a Nikon A1R inverted confocal microscope. Primary antibodies used include anti-APOL1 (5.17D12, 1:500, Genentech), anti-Calnexin (NB300518, 1:500, Novus), and anti-Tom20 Coralite 647 (CL647-11802, 1:500, Proteintech Group). Nuclei were stained with DAPI (1:10,000).

### Autophagy measurements

BMDMs were seeded onto 96-well plates and treated with 5 ng/mL of IFNγ and 50 µM of oxLDL for up to 72 hours. The autophagy assay kit (BioRad) was used to measure fluorescence. Cells were incubated with Autophagy Probe Red reagent for 1 hour. Cells were washed with cell culture media 3 times and resuspended in flow buffer (1X PBS, 0.5% BSA, 0.05% Sodium Azide). Cellular fluorescence was measured at 590 nm excitation and 620 nm emission with LSRFortessa Flow Analyzer (BD Biosciences).

### Thallium flux analysis

BMDMs were seeded into a 384-well tissue culture dish and treated with 5 ng/mL of IFNγ or PBS control for 8 hours. Thallium flux was measured using the Brilliant Thallium assay (ION Biosciences). BMDMs were incubated with a thallium-selective fluorescent indicator for 1 hour prior to assay. Then, 1 μM VX-147 (GLPBio, GC64339) or DMSO control was added to the cells for 30 minutes prior to the addition of thallium. Fluorescence was measured using the Panoptic kinetic imaging system (WaveFront Biosciences) and the slope of the fluorescence after thallium addition was quantified using WaveExplorer (WaveFront Biosciences). The data was a combination of 8 biological replicates over 2 separate experiments.

### Transmission electron microscopy

BMDMs and iPSDMs were cultured on Thermanox plastic coverslips (Thermo Fisher Scientific, 174950) and treated with 5 ng/mL of IFNγ or PBS control for 8 hours. Cells were then fixed with 0.1 M sodium cacodylate buffer (pH 7.35) with 2% paraformaldehyde and 2.5% glutaraldehyde. Cells were post-fixed with 2% osmium tetroxide and stained *en bloc* with 3% uranyl acetate. Samples were embedded in resin and sectioned into 70 nm thin sections onto copper grids. The samples were post-stained with 3% uranyl acetate and Reynolds lead citrate. Images were collected on an FEI Spirit Transmission Electron Microscope.

### Quantification of mitochondrial area and circularity

Mitochondria in BMDMs were analyzed as previously described(101). Briefly, mitochondria were outlined individually and measured in macrophages using ImageJ. Area and circularity were derived from the outlines of mitochondria in each sample. Circularity was calculated with the following equation: Circularity = (4 * π * Area) / (Perimeter^2)

### Seahorse metabolic flux analysis

Oxygen consumption rate (OCR) and extracellular acidification rate (ECAR) of macrophages were measured using a Seahorse XFe96 analyzer (Agilent). The cells were seeded into a 96-well assay plate at 2 × 10^5^ cells/well according to the Seahorse manual. The cells were maintained overnight at 37°C supplemented with 5% CO_2_. The next day, the media was replaced with Seahorse XF RPMI medium (Agilent, 103681-100) and the cells were equilibrated in a non-CO2 incubator at 37°C. Oxidative phosphorylation was measured with the Mito Stress Test (Agilent, 103015-100) as follows: OCR and ECAR measurements occurred after mixing the wells. Oligomycin (1 µM), FCCP (1.5 µM), and Rotenone/Antimycin A (0.5 µM) were injected sequentially into the cells with three measurement points in between each injection. Glycolytic rate was measured with Glycolytic Rate Assay (Agilent, 103344-100) as follows: ECAR and proton efflux rate (PER) were measured after Rotenone/Antimycin A (0.5 µM) and 2-deoxyglucose (50 mM) was injected sequentially into the cells with 3 measurement points between each injection. Data was normalized to cell concentration as measured by CyQuant assay (Invitrogen, C7026).

### Hydrophilic metabolite isolation and reconstitution

Intracellular hydrophilic metabolites were isolated and analyzed to quantify relative abundance. Metabolites were isolated from G0, G1, and G2 BMDMs on dry ice with 1 mL of 80% v/v methanol. The insoluble fraction was precipitated at -80°C overnight and the soluble fraction was dried down using SpeedVac vacuum concentration (Thermo Fisher Scientific). 60% acetonitrile was added to the tube for reconstitution followed by overtaxing for 30 sec. The sample solution was then centrifuged for 30 min @ 20,000g, 4 °C. Supernatant was collected for LCMS analysis.

### Metabolite profiling and analysis

Metabolite profiling was carried out as previously described(102). Samples were analyzed by High-Performance Liquid Chromatography and High-Resolution Mass Spectrometry and Tandem Mass Spectrometry (HPLC-MS/MS). Specifically, the system consisted of a Thermo Q-Exactive in line with an electrospray source and an Ultimate3000 (Thermo Fisher Scientific) series HPLC consisting of a binary pump, degasser, and auto-sampler outfitted with an Xbridge Amide column (Waters; dimensions of 3.0 mm × 100 mm and a 3.5 µm particle size). The mobile phase A contained 95% (vol/vol) water, 5% (vol/vol) acetonitrile, 10 mM ammonium hydroxide, 10 mM ammonium acetate, pH = 9.0; B was 100% Acetonitrile. The gradient was as follows: 0 min, 15% A; 2.5 min, 64% A; 12.4 min, 40% A; 12.5 min, 30% A; 12.5-14 min, 30% A; 14-21 min, 15% A with a flow rate of 150 μL/min. The capillary of the electrospray ionization (ESI) source was set to 275 °C, with sheath gas at 35 arbitrary units, auxiliary gas at 5 arbitrary units and the spray voltage at 4.0 kV. In positive/negative polarity switching mode, an m/z scan range from 60 to 900 was chosen and MS1 data was collected at a resolution of 70,000. The automatic gain control (AGC) target was set at 1 × 10^6^ and the maximum injection time was 200 ms. The targeted ions were subsequently fragmented, using the higher energy collisional dissociation (HCD) cell set to 30% normalized collision energy in MS2 at a resolution power of 17,500. Besides matching m/z, target metabolites are identified by matching either retention time with analytical standards and/or MS2 fragmentation pattern. Data acquisition and analysis were carried out by Xcalibur 4.1 software and Tracefinder 4.1 software, respectively (both from Thermo Fisher Scientific).

Analysis was performed using MetaboAnalyst 6.0 to determine the differential expression of metabolites(103). Metabolites were first TIC-normalized before uploading to the MetaboAnalyst web server. By default, missing values were replaced by 1/5 of the minimum positive value for each metabolite. Only auto-scaling was applied during the normalization step. For functional analyses, TIC-normalized abundances were used for quantitative enrichment analysis with the "Enrichment Analysis" and "Pathway Analysis" modules using the same parameters as above. Volcano plots and relative abundance of individual metabolites were derived from “Enrichment Analysis.”

### Cell viability assay

G0, G1, and G2 BMDMs were seeded into a 96-well plate and treated with 0-100 ng/mL of mouse IFNγ for 4, 8, and 24 hours. Cell viability was determined using CellTiter-Glo (Promega, G7570) according to the manufacturer’s protocol and measuring luminescence. Data was normalized to the untreated control wells.

### Sex as a biological variable

All BMDM experiments were completed with even male and female sex representation. Data findings were similar between both sexes. For iPSDM experiments, one male and one female line were used.

### Statistics

Tests for statistical significance were performed using GraphPad Prism software. Values are presented as mean ± SD except where noted. For comparisons between 2 groups, statistical differences were determined by a 2-tailed, unpaired Student’s t-test. For comparison between more than 2 groups, ANOVA with Tukey’s multiple-comparison test was used. P values less than 0.05 were considered statistically significant.

### Study approval

All animal experiments were approved by the Animal Care and Use Committee of Northwestern University. All experiments were performed in accordance with the relevant guidelines and regulations.

## Data availability

All data in this article will be shared by the lead contact upon request. Metabolomics input data are available in the supplement.

## Author Contribution

EL: Conception of work, data acquisition, data analysis, data interpretation, drafting of manuscript

MW: analytical design, data analysis, data interpretation, drafting of manuscript

AOK: data acquisition, data analysis, manuscript review

TC: data acquisition, data interpretation

JYY: data acquisition, manuscript review

VG: data acquisition, data analysis, manuscript review

AOD: data acquisition

SI: experimental design, manuscript review

NSC: data interpretation, manuscript review

HZ: experimental design, manuscript review

EBT: data interpretation, manuscript review

JL: Conception of work, data analysis, data interpretation, drafting of manuscript

All co-authors edited and approved the manuscript.

## Acknowledgements

The authors are grateful for the APOL1 antibodies received from Suzie Scales at Genentech. Research reported in this manuscript was supported by R01-DK131521-03, F31-DK131884-02. Research reported in this publication was also supported by the National Institute of Diabetes and Digestive and Kidney Diseases of the National Institutes of Health under Award Numbers U2CDK129917 and TL1DK132769. Metabolomics services were performed by the Metabolomics Core Facility at Robert H. Lurie Comprehensive Cancer Center (generously supported by NCI CCSG P30 CA060553) of Northwestern University. Confocal imaging and electron microscopy work was performed at the Northwestern University Center for Advanced Microscopy (RRID: SCR_020996) generously supported by NCI CCSG P30 CA060553 awarded to the Robert H Lurie Comprehensive Cancer Center. Flow cytometry work was done at the Northwestern University RHLCCC Flow Cytometry Facility and was supported by the Cancer Center Support Grant (NCI CA060553). Metabolomics experiments and Seahorse assays were performed at the Metabolomics Core Facility at Robert H. Lurie Comprehensive Cancer Center (generously supported by NCI CCSG P30 CA060553) of Northwestern University. The content is solely the responsibility of the authors and does not necessarily represent the official views of the National Institutes of Health.

---

**Supplemental Figure 1.**
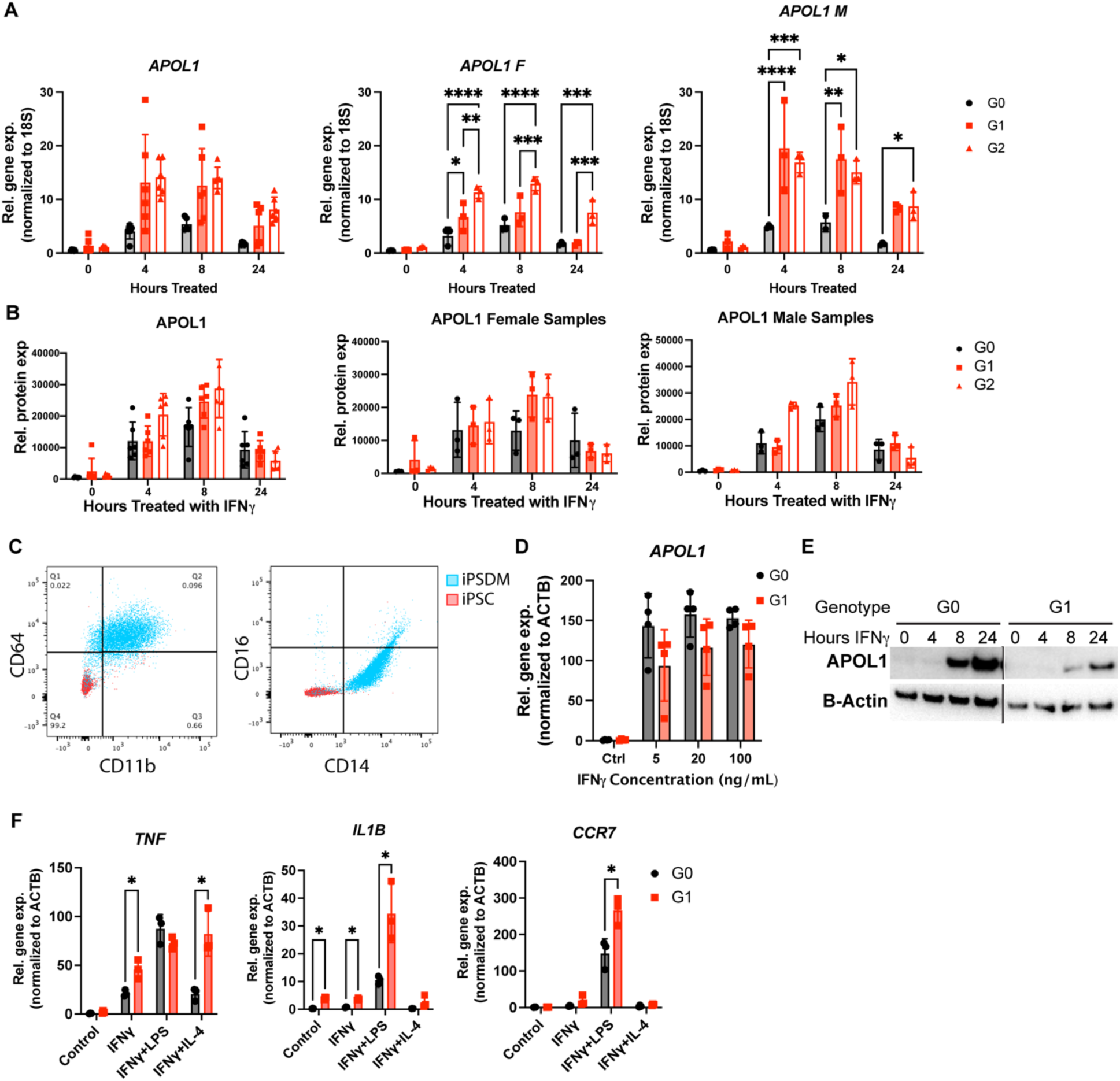
A**P**OL1 **expression in BMDMs and iPSDM is increased with IFNγ (A)** Gene expression of *APOL1* measured with qRT-PCR in combined sex, female, and male BMDMs. **(B)** Quantification of APOL1 protein expressed measured with western blot in combined, female, and male BMDMs. **(C)** Flow cytometric analysis of CD11b, CD64, CD14, and CD16 surface expression on G1 iPSDM and G1 iPSCs. **(D)** Gene expression of *APOL1* measured in G0 and G1 iPSDMs treated with IFNγ. **(E)** western blot of APOL1 in G0 and G1 iPSDMs treated with IFNγ. BMDM experiments were performed in BMDMs from 3-6 mice per genotype per group, with both sexes represented. **(F)** Gene expression of *TNA, IL1B,* and *CCR7* measured with qRT-PCR in G0 and G1 iPSDMs treated with IFNγ (25 ng/mL), LPS (10 pg/mL), and IL-4 (10 ng/mL) for 24 hours. BMDM experiments in **(A, B)** were completed in BMDMs from 4 mice from each genotype and each sex. iPSDM experiments in **(C, D)** were performed in 2 separate differentiations across 2 independent experiments. Experiments in **(E)** were performed with 3 separate wells of iPSDM. Data are expressed as mean ± SD. *p<0.05, **p<0.01 ***p<0.005 ****p>0.001. 2-way ANOVA with Tukey’s Multiple Comparison Test

**Supplemental Figure 2.**
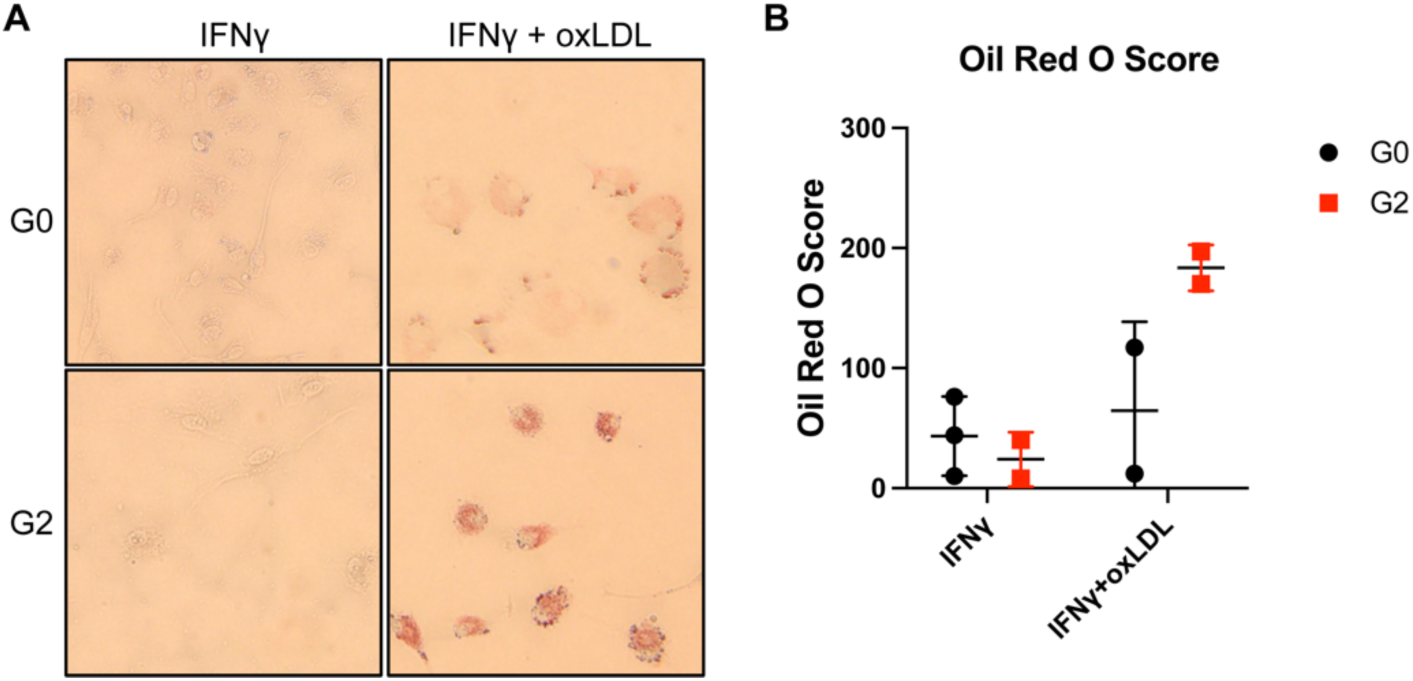
G**2 BMDMs from the C57Bl/6J background increase lipid accumulation with oxLDL incubation. (A)** Oil Red O (ORO) stained images of BMDMs in the presence or absence of 5ng/mL IFNγ and 50 μg/mL oxidized LDL (oxLDL) for 72 hours. BMDMs were generated from G0 and G2 mice from the C57Bl/6J background**. (B)** Quantification of the Oil Red O stain. Experiments were completed in BMDMs from 2-3 mice per genotype per group, with both sexes represented. Data are expressed as mean ± SD.

**Supplemental Figure 3.**
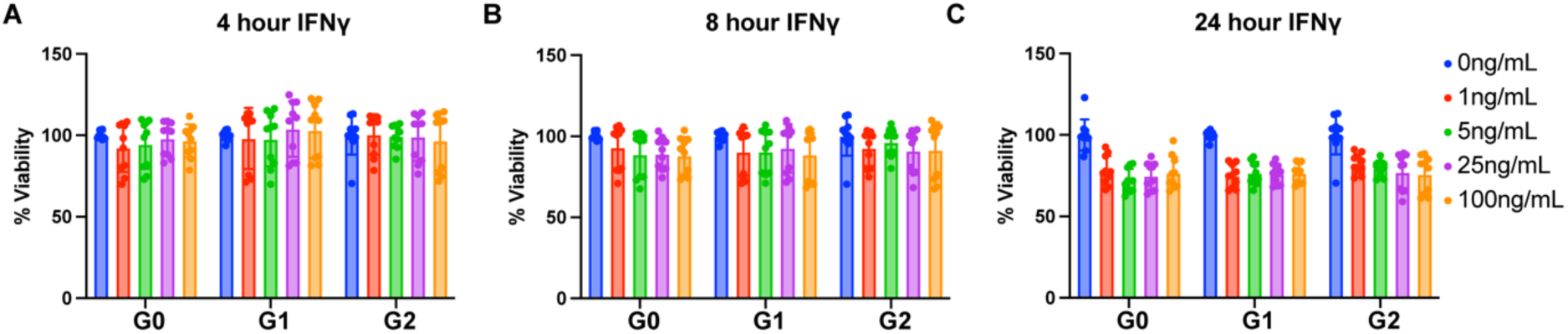
C**e**ll **viability of BMDMs treated with IFNγ is slightly decreased at 24 hours (A-C)** Cell viability of BMDMs treated with 0-100 ng/mL IFNγ for 4, 8, and 24 hours measured with CellTiter-Glo. Experiments were performed using BMDMs from 8 mice across 2 independent experiments, with both sexes represented. Data are expressed as mean ± SD.

**Supplemental Figure 4.**
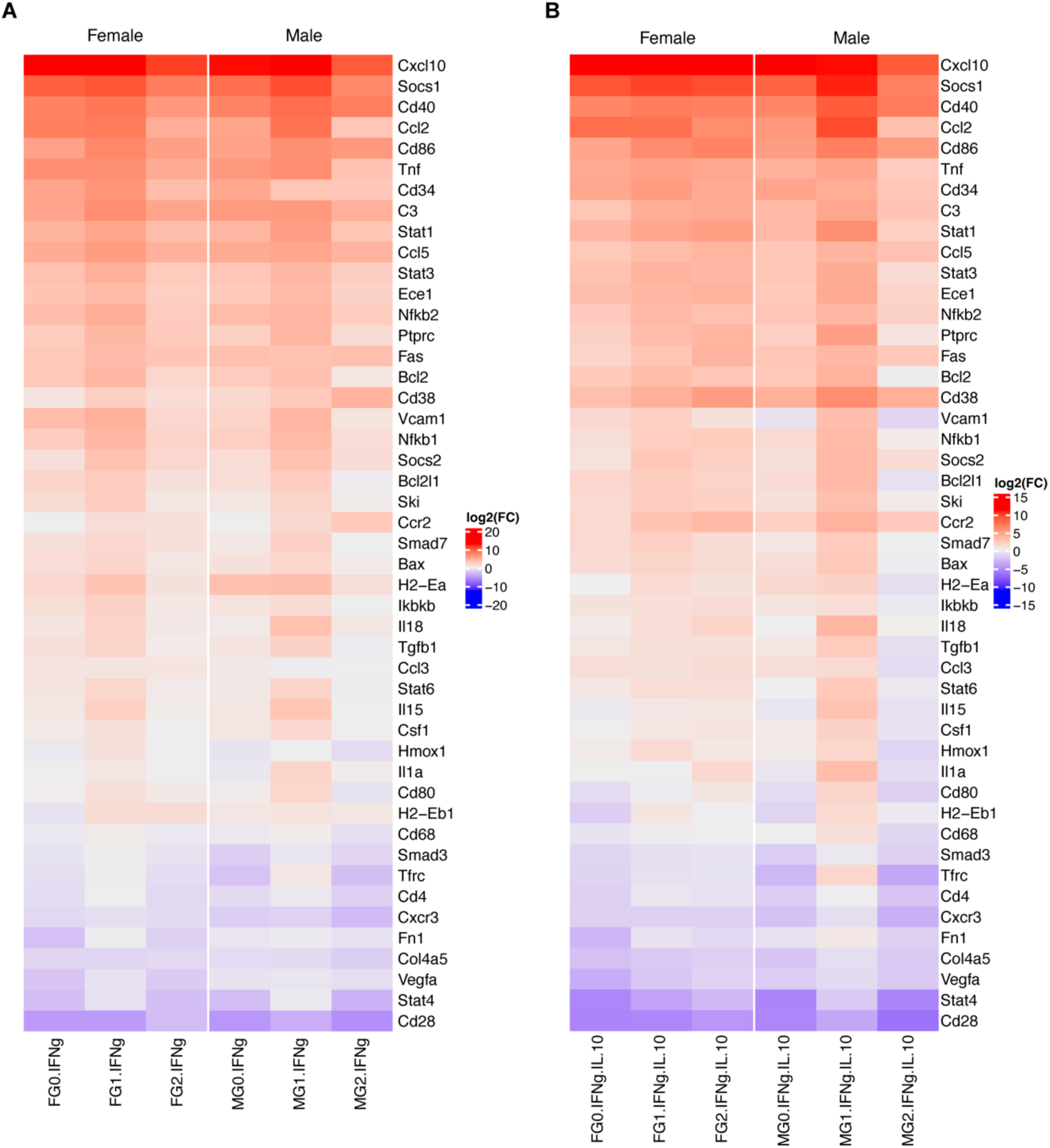
q**P**CR **array of immune genes in female and male BMDMs. (A-B)** qPCR array of immune-related genes in female and male BMDMs treated with IFNγ (5 ng/mL) and IL-10 (10 ng/mL). Experiment was completed in BMDMs from 2 mice per genotype per group.

**Supplemental Figure 5.**
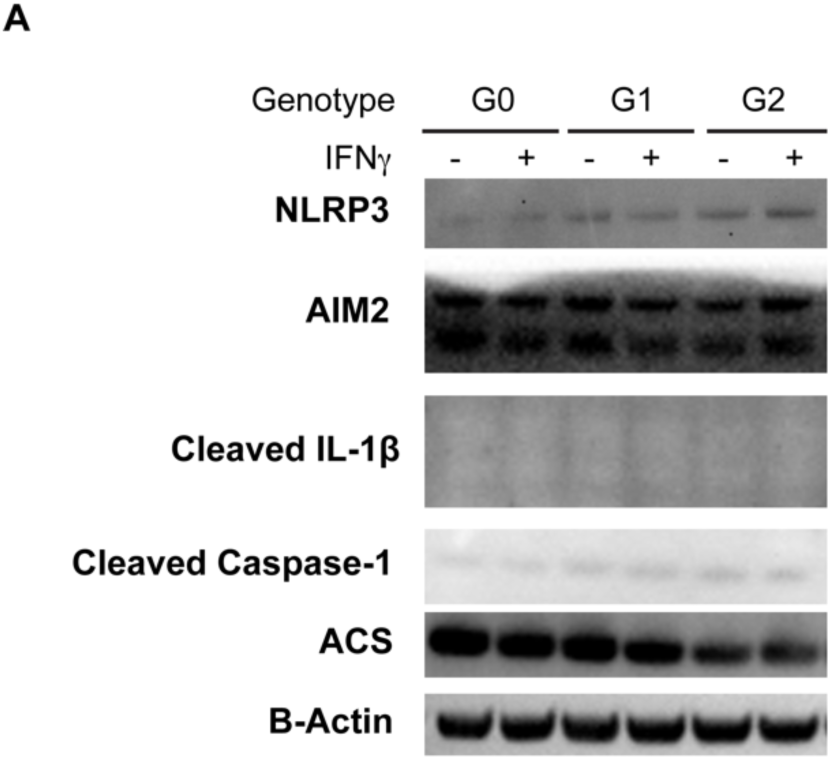
N**L**RP3 **inflammasome proteins are not activated with IFNγ treatment in APOL1 BMDMs. (A)** Representative western blot of NLRP3, AIM2, cleaved IL-1β, cleaved Caspase-1, and ACS in G0, G1 and G2 BMDMs treated with IFNγ (5 ng/mL) or control. Experiments were performed in BMDMs from 4 mice per genotype per group, with both sexes represented.

**Supplemental Figure 6.**
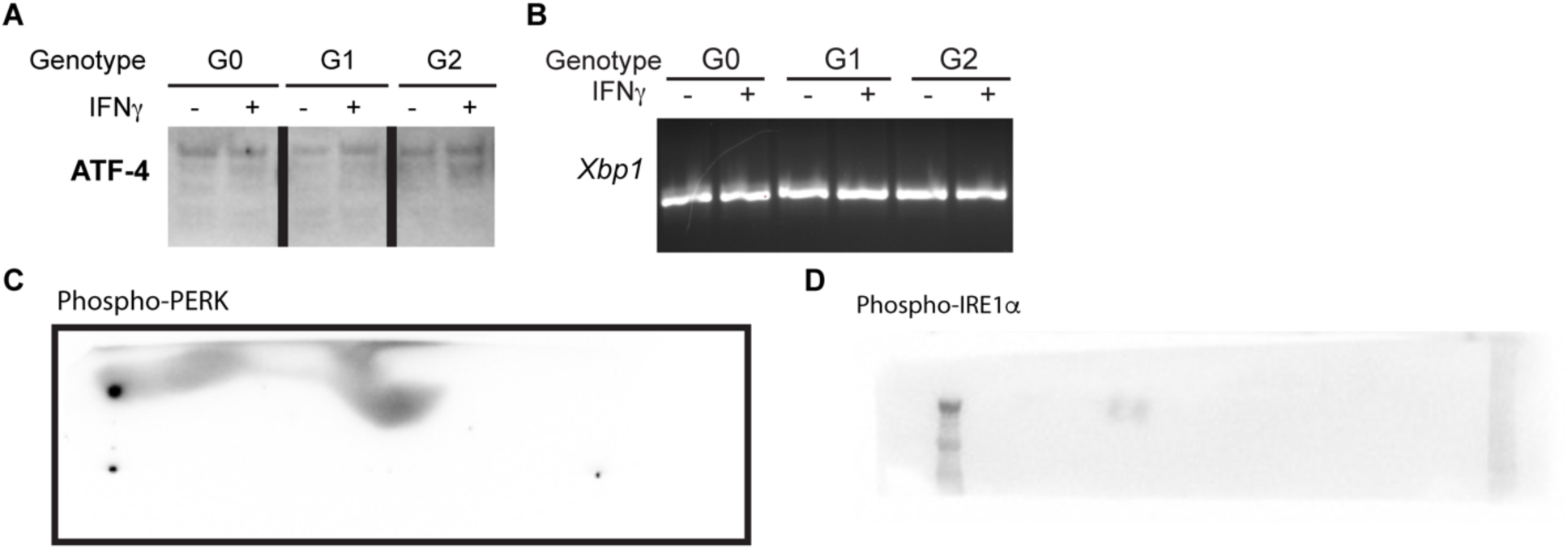
E**R stress proteins and genes are not expressed in APOL1 BMDMs with IFNγ treatment.** Representative western blot of ATF4 **(A)**, phospho-PERK **(C)**, phospho-IRE1α **(D)** and RT-PCR of Spliced *Xbp1* **(B)** in G0, G1 and G2 BMDMs treated with IFNγ (5 ng/mL). Experiments were completed in BMDMs from 4 mice per genotype per group, with both sexes represented.

**Supplemental Figure 7.**
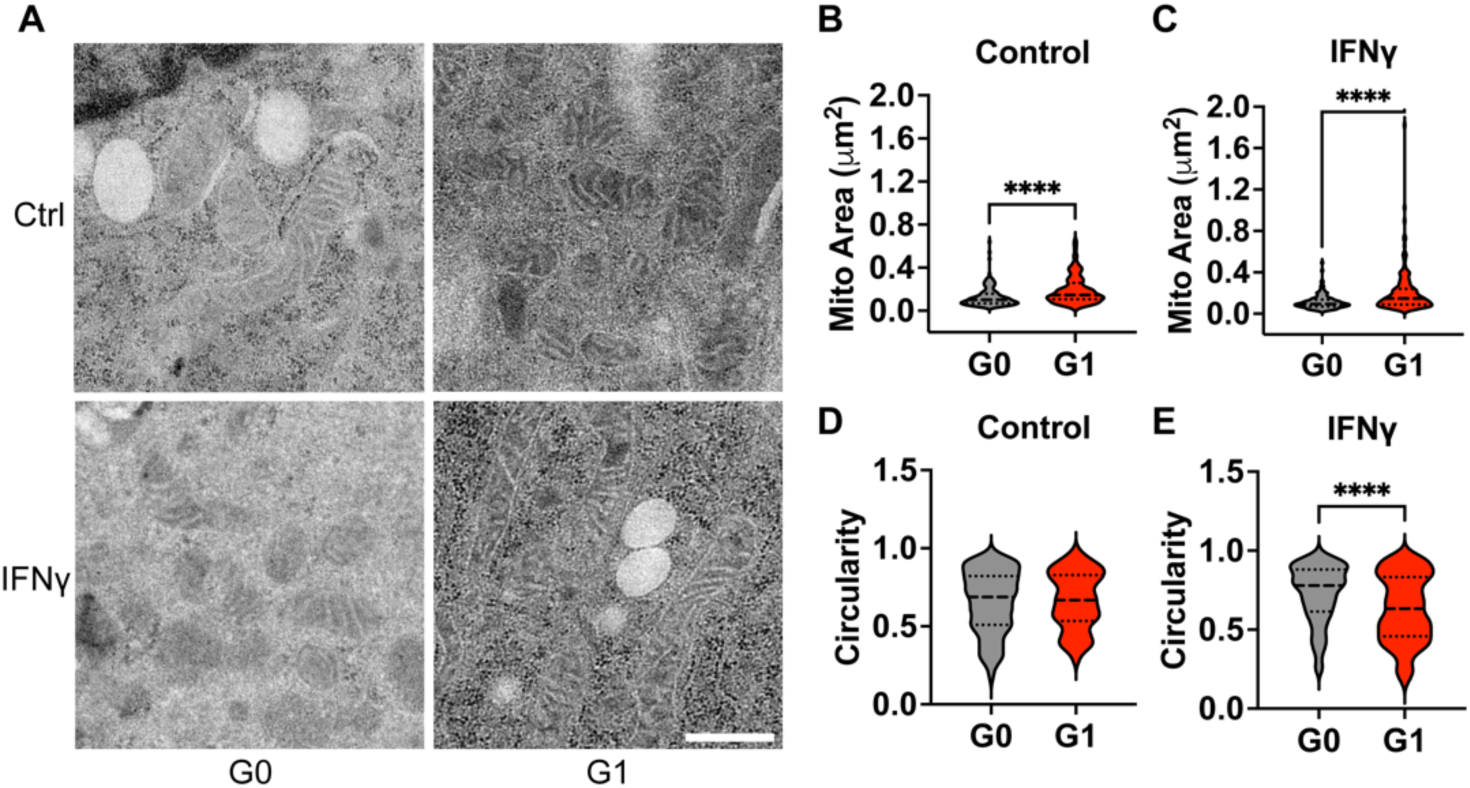
M**i**tochondrial **area is increased in G1 iPSDM compared to G0** G0 and G1 iPSDMs were treated with 20 ng/mL IFNγ for 8 hours.TEM images were collected of at least 20 cells and over 100 mitochondria were quantified for area and circularity. **(A)** Representative images of mitochondria in control and treated iPSDM. **(B, C)** Quantification of mitochondrial area in iPSDM. **(D, E)** Quantification of mitochondrial circularity in iPSDM. Experiments were performed in iPSDM from 4 separate differentiations across 2 independent experiments. Data are expressed as median and interquartile range. Unpaired *t* test. ****p <0.001

**Supplemental Figure 8.**
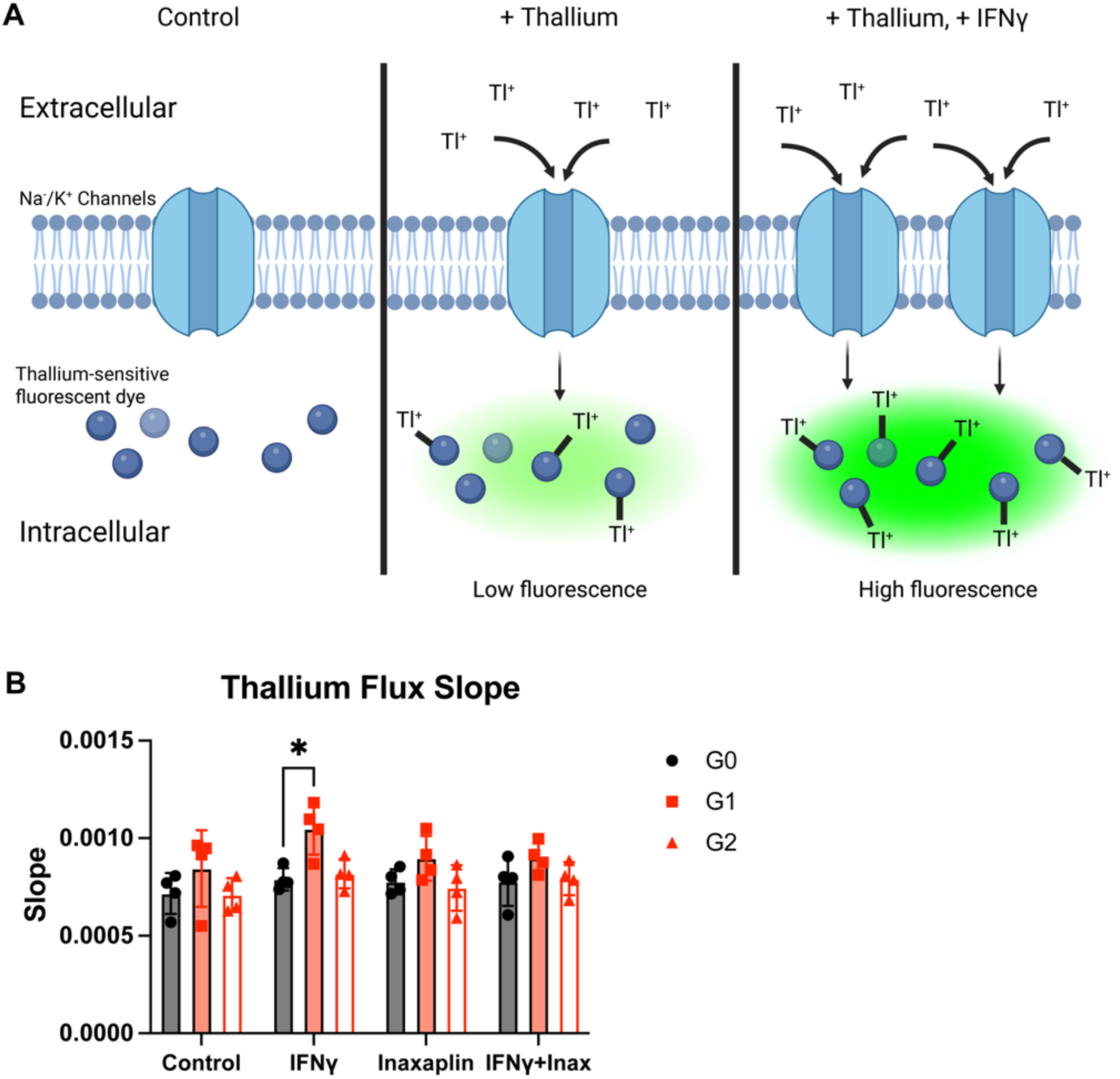
T**h**allium **flux in APOL1 BMDMs is unchanged with IFNγ and VX-147 treatment. (A)** Diagram of Thallium flux assay. **(B)** Slope of fluorescence after thallium addition to G0, G1 and G2 BMDMs treated with IFNγ (5 ng/mL), VX-147(1 μM) or DMSO control. Experiments were performed in BMDMs from 4 mice per genotype per group, with both sexes represented. Data are expressed as mean ± SD. Unpaired *t* test. *p <0.05

## Notes

The authors have declared no conflict of interest exists.

### Competing Interest Statement

The authors have declared no competing interest.

